# A Fundamental Approach to Buoyant Density Determination by DGE-AUC

**DOI:** 10.1101/2025.04.28.650698

**Authors:** Alexander E. Yarawsky, Paola Cardenas Lopez, Johannes Walter, Michael T. DeLion, Lake N. Paul

## Abstract

Density gradient equilibrium analytical ultracentrifugation (DGE-AUC) was introduced in 1957 by Meselson, Stahl, and Vinograd. The method saw significant use for nucleic acids, proteins, synthetic polymers, and viruses throughout the 1960s and 1970s. Since then, DGE-AUC has seen continued use in the polymer/nanoparticle field and the genomic DNA field. New developments in medicine have revived interest in the technique for the characterization of cell and gene therapeutics. While several theoretical (“*model*-*dependent*”) approaches exist to determine density at any given point along a density gradient at equilibrium, there is ample evidence in the 50+ years of density gradient literature that indicates the presence of pressure effects, solvent compressibility, and general nonideal behavior of the gradient medium that are not easily accounted for in models describing the density gradient. These complications led to the requirement for using reference standards at standard conditions where empirical relationships have been developed and tested over many years. With an interest in buoyant density determination for virtually any particle of various composition, an approach that does not rely on reference standards is desirable. The current manuscript details a fundamental “*model*-*independent*” method for determination of the buoyant density of a particle via DGE-AUC. An examination of this novel DGE-AUC method is presented in the context of NISTmAb and lambda bacteriophage DNA in a CsCl gradient, as well as polystyrene beads in a sucrose gradient. The method described herein is broadly applicable as a model-independent approach to determining the buoyant density of a particle in a density gradient medium.

**Highlights:**

- Density gradients are a classical approach to separating particles.
- Analogous experiments can be performed in an analytical ultracentrifuge.
- Historical methods exist but require assumptions about the density gradients.
- A robust and flexible model-independent approach is presented.
- Several applications of the new approach are demonstrated.

**Graphical Abstract:** 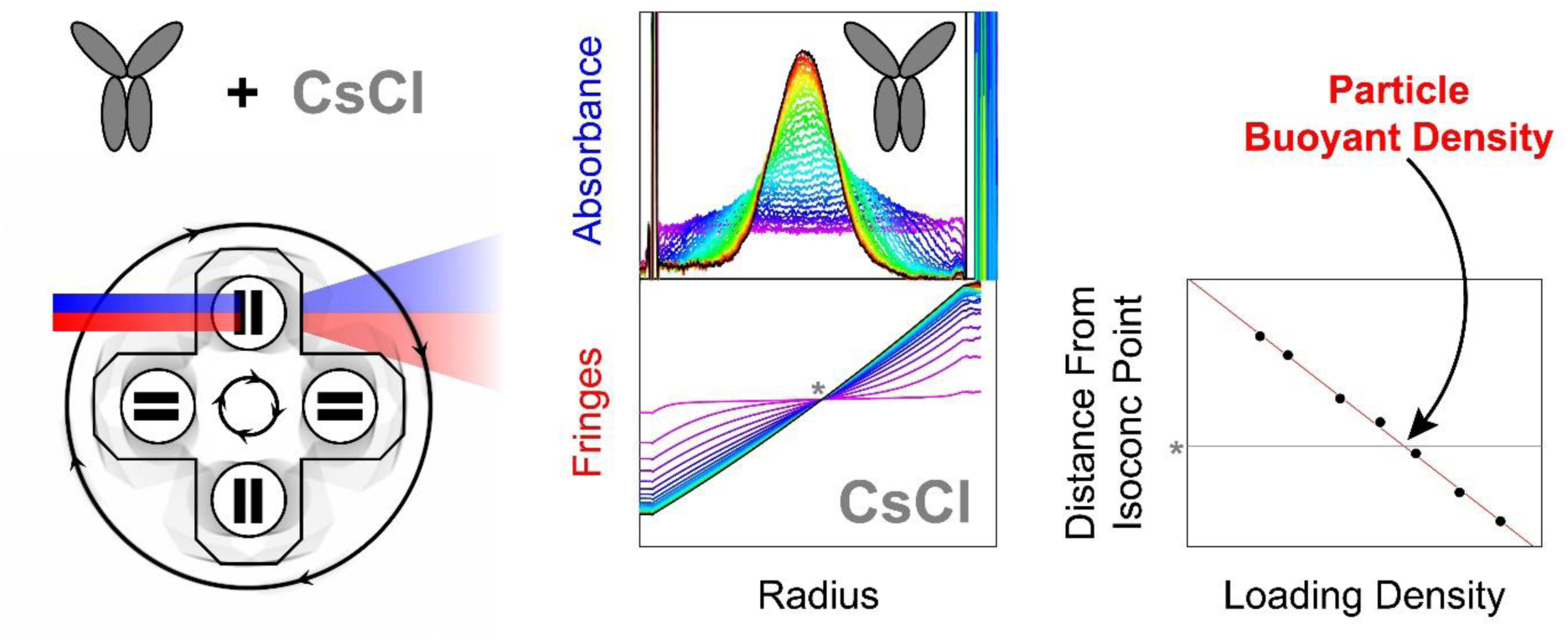

## 1. Introduction

Analytical ultracentrifugation (AUC) has seen a resurgence of interest recently, primarily due to the need to characterize and quantitate viral capsid loading states for adeno-associated virus (AAV) therapeutics. The commonly used sedimentation velocity (SV-AUC) experiments are of particular importance for their ability to provide high resolving power for not only empty and full capsids, but also partially filled capsids. This has placed SV-AUC as the gold-standard analytical technique in the field [1–5], which has, in part, led to development of cGMP (current good manufacturing practices) compliant analysis software [6–8]. In addition to SV-AUC, other classical AUC methods including analytical band AUC [9–11] and density gradient equilibrium AUC (DGE-AUC) have become popular [10,12–15], as they have lower sample requirements than SV-AUC. This manuscript focuses on the development of a novel, fundamental approach to DGE-AUC for accurate buoyant density determination of a particle.

Meselson et al. first presented the density gradient equilibrium AUC experiment in 1957. The experiment involved the addition of density gradient forming material (i.e., CsCl) to an analyte (i.e., DNA). Upon centrifugation, CsCl forms a concentration gradient, which in turn forms a density gradient. The DNA will sediment or float based on its intrinsic buoyant density and the density of solution in its local environment. This leads to the banding of DNA at a given position along the density gradient [16]. Much of the early work on understanding density gradients in the ultracentrifuge came from Jerome Vinograd’s laboratory in the late 1950s and early 1960s [16–20]. This work aimed to understand the behavior of DNA and viruses in a density gradient formed by CsCl or other salts. It laid out the fundamental theory involved and indicated that solvent compressibility and other pressure effects were experimentally present [18,19]. From the late 1950s to the present day, laboratories interested in genomic DNA characterization have been using the approaches first laid out by Vinograd and colleagues [16,17,21]. It was determined by several parties concomitantly that a linear relationship existed between the guanine-cytosine (GC) content of DNA and the buoyant density in CsCl [22–24]. The use of CsCl-based DGE-AUC experiments led to many important discoveries which predated gene sequencing and the eventual sequence of the first human genome [21]. Even with current, easily accessible sequencing techniques, there is still a valuable place for CsCl DGE-AUC experiments in obtaining rapid and reliable information on the composition and heterogeneity of complete genomes [21,25]. Very early on, a reference marker of known buoyant density – *E. coli* DNA or bacteriophage DNA – began to be incorporated into DGE-AUC experiments. This allowed for a simpler determination of the buoyant density of an unknown DNA sample via a relationship between a radial distance in the AUC cell and the CsCl density increment at a given rotor speed [21,24,26,27]. Inclusion of a reference standard was said to be: “By far the most satisfactory way of determining buoyant density…” [27]. According to a review of the technique in 1973, differences in buoyant density down to 0.0005 g/cm^3^ could be determined significantly, while absolute buoyant densities were typically less accurate [27]. Clearly, there is a rich history of using DGE-AUC for DNA characterization in an analytical context.

The usefulness of DGE-AUC certainly extends beyond DNA. Density gradients in a preparative centrifuge are a classical approach to the purification of viruses [28,29]. Naturally, analytical methods followed. Berkowitz and Philo [30], in 2007, demonstrated the use of CsCl-based DGE-AUC in a modern AUC to evaluate the purity of adenovirus preparations. More recently, after an explosion of therapeutic interest in adeno-associated viruses (AAV), DGE-AUC experiments once again resurfaced. Several studies clearly demonstrate the utility of DGE-AUC for evaluation of vector homogeneity in drug delivery systems [10,12–15]. In the 1960s and 1970s, Ifft and colleagues pursued the use of DGE-AUC for protein characterization – specifically in regards to solvation and surface charges via pH-dependent buoyancy titrations [31,32].

Further expansion of the utility of DGE-AUC was demonstrated by Hermans and Ende, who presented the application to polymers [33]. From the 1980s through the early 2000s, Mächtle developed density gradient approaches with a strong focus on polymer characterization [34–36]. Others have also demonstrated interest and applications of density gradients for nanoparticles [37,38]. Because a particle’s buoyant density is an intrinsic property based on its composition and surface chemistry, there could be virtually endless applications.

With the foundational DGE-AUC work being performed on the Spinco Model E analytical ultracentrifuge, we now explore advantages provided by the interferometry optical system (for measuring refractive index differences between the two AUC centerpiece channels) implemented on the Beckman Optima XL-I or ProteomeLab XL-I released in the 1990s and early 2000s, as well as the most recent Optima AUC. Rossmanity and Mächtle proposed using these modern interference optics. While shallow density gradients of water/metrizamide mixtures provided suitable results, they observed that steeper density gradients are problematic in the calculation of mass conservation used for converting the interference data to an absolute refractive index – used finally to obtain the density across the gradient. Thus they recommended using calibration marker particles and the Schlieren optics available only on the Spinco Model E AUC released in 1947 [35,36].

Ifft and colleagues were first to introduce the term isoconcentration point in 1961 [17]. This is the radius where the final density at equilibrium is equal to the starting density. Model-dependent analyses have taken advantage of this point – estimated as the root-mean-square position between the solution column meniscus and bottom, from which the density along the gradient is extrapolated.

In this current manuscript, a novel approach to determine the buoyant density of a particle from DGE-AUC experiments is described. This approach aims to be rigorous and fundamental, with little to no assumptions required about the system composition (the particle or the density gradient medium). It is a global analysis that allows for robust evaluation of data and error estimation of any relevant parameters. Testing focused on the NIST monoclonal antibody reference material (NISTmAb; RM 8671), λ DNA, and polystyrene (PS) beads. However, the method outlined herein should be applicable to nearly any particle of interest. Furthermore, heterogeneous particles and multi-component particles, such as AAV with different loading states, mixtures of protein and DNA components, multi-component virus-like particles (VLP), polymers, and nanoparticles may all be characterized using this approach. While CsCl and sucrose were used here, various other density gradient media are available.

## 2. Experimental

### 2.1 Materials

Monoclonal antibody (NISTmAb; Reference Material^®^ 8671; Lot 14HB-D-001) was obtained from the National Institute of Standards and Technology (NIST). λ DNA was obtained from NEB (N3011S). Styrene (≥99%, Reagent Plus), acrylic acid (99%, 200 ppm MEHQ Inhibitor), ammonium persulfate (≥98%), and sodium dodecyl sulfate (SDS, ≥98%, Reagent Plus) were purchased from Sigma-Aldrich. Sodium hydroxide (≥98%, Ph. Eur., USP, BP, in pellets), aluminum oxide (90 neutral), and D(+) saccharose (≥99%) were purchased from Carl Roth GmbH. For water, a Purelabflex (Elga Lab water) purification unit was used (18.2 MΩ cm). Styrene was purified to remove the inhibitor by extraction with a 10 wt.% aqueous sodium hydroxide solution and by passing through an aluminum oxide column. The cleaned styrene was stored at 8°C for a maximum of 2 months before use. Optical grade CsCl (C3139) was obtained from MilliporeSigma and prepared in water or 20 mM Tris pH 8.0 (Teknova T1080).

NISTmAb was dialyzed into a 50 mM Tris pH 8.0 solution prior to DGE-AUC. NISTmAb was stored at-80°C in aliquots, with a fresh aliquot used for each experiment.

SigmaPlot (v12.5) was used for producing figures and fitting data.

### 2.2 Synthesis of colloidal PS particles

PS beads were synthesized by surfactant-free emulsion polymerization [39]. Briefly, 990 mL of ultrapure water was heated to 76°C in a 2 L three-necked-flask equipped with a reflux condenser under stirring at 500 rpm and constant purging with nitrogen. Next, 28 g of styrene was added. Subsequently, 0.4 g of acrylic acid and 0.4 g of ammonium persulfate, both dissolved in 5 mL of ultrapure water, were added in 10-minute intervals. The addition of the initiator marked the start of the reaction. After 22 hours, the nitrogen flow and heating were stopped, the colloid was left to cool, and the solution was subsequently filtered using a lint-free wipe (KIM Wipe, Kimberly-Clark). After synthesis, the mean particle diameter of the PS beads was measured using scanning electron microscopy and determined to be x_0.5,SEM_ = (247.3 ± 2) nm. For DGE-AUC experiments, PS particles were transferred to water-based sucrose solutions containing 1 mM SDS.

### 2.3 Densitometry (NISTmAb and DNA in CsCl)

Density was measured using an Anton Paar DMA5000 at 20°C. An air-water density adjustment was performed using the current air pressure according to www.weather.gov and pure water density standard (DENWAT3 from MilliporeSigma). A density check was then performed on water to ensure proper adjustment. Between samples, approximately 5 mL of water was used to wash the instrument three times, followed by a similar procedure using 70% isopropyl alcohol. The DMA5000 was then allowed to dry using the built-in air pump for 3-5 minutes. The water density standard was run periodically to ensure the instrument did not drift and to ensure adequate cleaning between samples.

The concentration of NISTmAb was measured in a NanoDrop One (ThermoFisher) UV/Vis using a 1 cm microcuvette based on absorbance at 280 nm with a baseline correction at 400 nm.

### 2.4 Densitometry (PS in sucrose)

The density of sucrose solutions was determined using an Anton Paar DMA5000 density meter. Prior to the measurements, air and water checks with a pure water density standard (VWR) were performed. The oscillating tube was cleaned after each measurement with ultrapure water, pure ethanol and ultrapure water rinsing cycles. To determine the most suitable concentration of PS particles for DGE-AUC experiments, the UV/Vis spectra of PS dispersions were taken using a Lambda 35 spectrometer (PerkinElmer). The evaluated wavelength range was 350 nm to 650 nm with a spectral resolution of 1 nm using a cuvette with a 1 cm optical path length.

### 2.5 DGE-AUC data collection (NISTmAb and DNA in CsCl)

NISTmAb and DNA experiments were conducted using a Beckman Coulter Optima XL-I with an 8-hole rotor, except when collecting data above 50,000 rpm, where a 4-hole rotor was used. Experiments were performed at 20°C or 25°C. Aluminum centerpieces (Beckman Coulter) with a 12 mm pathlength were used along with sapphire windows.

Approximately 420 µL of sample (NISTmAb, CsCl, and 20 mM Tris pH 8.0) and reference solution (20 mM Tris pH 8.0) were loaded into the respective sectors unless otherwise specified in the text. Absorbance scans were collected with a radial resolution of 0.003 cm and in continuous mode at a wavelength of 230 nm (except where otherwise specified). A final scan after equilibrium was collected with 5 replicates with a radial resolution of 0.001 cm and in step mode. Interference data were also collected.

There was no time delay between scans or delay time before collection started. A radial calibration was performed for each detection mode prior to each experiment. Datasets were examined for equilibrium using the approach to equilibrium function in SEDFIT (v17.0).

### 2.6 DGE-AUC data collection (PS in Sucrose)

PS experiments were conducted using a Beckman Coulter Optima AUC equipped with an absorbance and interference detector and an 8-hole rotor. The samples were measured in titanium centerpieces (Nanolytics Instruments) with a 3 mm pathlength and sapphire windows. In all reference sectors, 110 µL of 1 mM SDS solutions were introduced. In sample sectors, 110 µL of PS beads in sucrose solutions at 1 mM SDS were given. All experiments were carried out at 20°C and a rotor revolution of 40,000 rpm. Absorbance scan settings were collected at a wavelength of 400 nm with a radial resolution of 0.002 cm. To observe the progress of the measurements, scans were taken every hour for a maximal duration of 96 hours. Interference scans were collected every 230 seconds, recording 1500 scans in total.

### 2.7 DGE-AUC data analysis

The final absorbance scan at equilibrium was loaded into Microsoft Excel (Version 2411), where a linear or polynomial baseline correction was performed manually. A data region was then fitted to a Gaussian distribution with the aid of the Solver Add-in using the following equation:

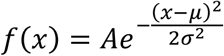

where *A* is the amplitude or signal intensity of the center of the distribution, μ is the mean or center of the distribution in terms radius, and σ is the standard deviation or width of the distribution in terms of radius. The Solver Add-in was used to minimize the sum of differences squared to determine the best fit using the GRG Nonlinear solving method. Options for the solving method were a Constraint Precision of 0.000001, 1% integer optimality, convergence of 0.0001, and forward derivatives. Alternatively, SigmaPlot (v12.5) was used for fitting the baseline-corrected data to a 3 parameter Gaussian to calculate 95% confidence intervals.

### 2.8 Interference data noise removal

Interference data contain two types of systematic noise. These are referred to as time-invariant noise (TI noise) and radially-invariant noise (RI noise). TI noise stems from debris or grease on cell windows and other components of the optical system and doesn’t change over time. The removal of TI noise is not typically required for this DGE-AUC method, however, scans of an empty AUC cell at the desired speed prior to loading them provides a simple approach to measuring TI noise. The scans can then be subtracted from the final scans at equilibrium. The RI noise, however, is important to remove from the interference data. RI noise is rooted in “mechanical and thermal variations caused predominantly by cycling of the Peltier heating and cooling system.” [40]. This results in fringe jumps, where the entire scan will be offset by an integer amount. Interference data were preprocessed in SEDVIEW [41] (v1.1.0 2007 available from http://www.rasmb.org/software/windows/sedview-hayes/). A data directory was chosen, and an interference data channel was selected. Once loaded, the jitter adjust position was set to a radius in the air-air space above the menisci. The signal in this region is assumed to be constant. Next, the region near the isoconcentration point was selected for integral fringe position (i.e., fringe jump removal), which multiplies or divides the scans by an integer such that all scans have a similar number of total fringes at the selected radius. The robustness of this can be seen in Fig. S1, where several different air-air regions were chosen for the jitter adjust along with several different radii for the integral fringe alignment. The scans were exported to Excel format and plotted in SigmaPlot (v12.5). Other programs utilize similar methods of RI noise removal or fit the RI noise as an additional model parameter. SEDVIEW was chosen due to its instantaneous re-calculating of the RI noise after adjusting the fringe jump removal radius and for its ease of use when handling many datasets from an experiment.

### 2.9 Hermans-Ende analysis

The Hermans-Ende approach has been mathematically described in detail elsewhere [33,36]. The gradient in the cell is formed due to the equilibrium of sedimentation and diffusion of the gradient forming material as described by Lamm’s equation in infinite time. In the following, we present the equation to supplementary calculate the solution local density *ρ_s_(r)*, in a cell at equilibrium conditions:

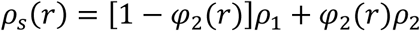

Where *r* is the radial position for evaluation, ρ_1_ is the density of the low-density material, typically a solvent, ρ_2_ is the density of the gradient forming material and φ_2_(*r*) is the local volume fraction of the gradient forming material defined as:

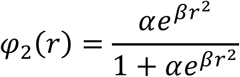

with:

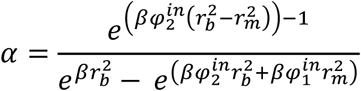

and:

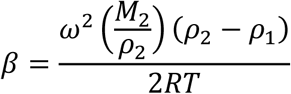

Where *φ_1_^in^* and *φ_2_^in^* are the initial volume fractions of solvent and gradient forming materials, respectively. *r_m_* and *r_b_* denote meniscus and bottom positions, ω is the angular velocity, *M*_2_ is the molar mass of the gradient forming material, *R* is the universal gas constant and *T* is the temperature.

Since the hydrostatic pressure has an effect on the built gradient [42], a pressure correction term is added to *ρ_s_(r)* as follows:

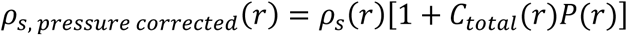

with:

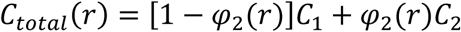

and:

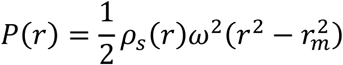

Where *C_total_(r)* is the local compressibility, *C*_1_ and *C*_2_ are the mixture components’ compressibilities and *P(r)* is the local hydrostatic pressure calculated with the uncorrected local density.

The Hermans-Ende equation enables the modeling of gradients and the prediction of local densities when the concentration of the gradient forming material is known.

Alternatively, if an estimate of the buoyant density of a particle is available, the equation can be used to determine the concentration of gradient forming material required to yield the particle at a particular radial position – something that is valuable for experimental design. However, the accuracy of these calculations depends on precise knowledge of all experimental parameters, making them susceptible to error.

Furthermore, as Mächtle and Börger opined, the Hermans-Ende theory may fail to accurately describe the density gradient within the cell when the mass percentage of the gradient forming material is too high [43]. For comparison, the Hermans-Ende equation was applied in this manuscript to analyze the buoyant densities obtained for PS beads. All parameter values used in the calculation are listed in Table S1.

### 2.10 Molecular weight determination

To estimate the apparent molecular weight *M_app_* of NISTmAb, the following equation from Schneider and Edelhoch [44] was used:

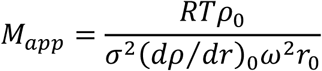

where *R* is the universal gas constant, *T* is temperature in Kelvin, ρ_0_ is the buoyant density of the particle (g/cm^3^), σ is the standard deviation of the Gaussian fit, (*dρ/dr*)_0_ is the density gradient at the peak’s radial center (*r*_0_), and ω is the angular velocity. The density gradient parameter was taken from the slope of the linear regression of a plot of loading density vs distance from isoconcentration point from a single experiment of NISTmAb in CsCl at 7 loading densities. This slope was multiplied by-1 to obtain a positive density gradient value – the density increases as the radius increases. The 95% confidence interval of the density gradient induced the largest error on the determination of apparent molecular weight, so this interval was used in calculations with the best fit values from the Gaussian peak fitting.

## 3. Results and discussion

### 3.1 Behavior of NISTmAb in CsCl solutions

To illustrate the fundamental principle of a DGE-AUC experiment, NISTmAb was added to three different concentrations of CsCl. Figure 1 shows the data collected during the centrifugation of the three samples. When a low concentration of CsCl was present, NISTmAb sedimented – much like a typical SV-AUC experiment (Fig. 1A). When a high concentration of CsCl was present, floatation was observed, where the protein moves toward the meniscus and the center of the rotor (Fig. 1B). These two observations illustrate that the buoyant density of the particle dictates whether it will sediment or float in a given solution density. When the particle is more dense than the solution, sedimentation occurs. When the particle is less dense than the solution, floatation occurs. Lastly, when the solution density is similar to the particle buoyant density, a “band” or peak is formed within the AUC cell (Fig. 1C). Specifically, the peak forms at the location along the density gradient at the radius where the solution density exactly equals the particle buoyant density. The system eventually reaches equilibrium, where no net movement occurs.

**Figure 1.**
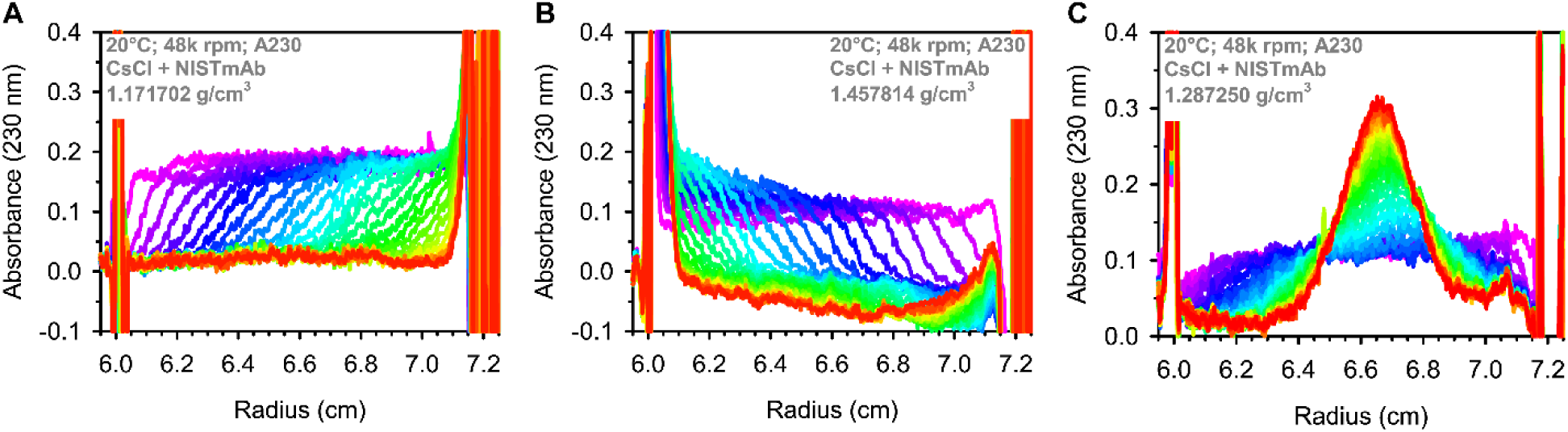
The impact of CsCl on NISTmAb sedimentation behavior. NISTmAb was examined in the presence of widely varying CsCl concentrations, which yielded very different solution densities. (A) NISTmAb in a solution density much lower than the buoyant density of NISTmAb, resulting in sedimentation. (B) NISTmAb in a high solution density, resulting in floatation. (C) NISTmAb in a solution density that is similar to its buoyant density, resulting in a band or peak near ∼6.65 cm at equilibrium. A total of 30 scans are shown in each dataset, where the rainbow color scheme reflects the time of the scan (magenta = first scan; red = last scan). Each panel shows the relevant sample and instrument parameters.

### 3.2 Monitoring Density Gradient Formation via interference optics

In the datasets in Figure 1, only part of the system is observable – the NISTmAb absorbance. To obtain a complete picture of the system, the interference optical system (i.e., Rayleigh interferometer or IF) may be used. The IF signal indicates the differences in the refractive index between a sample solution and reference solution – each loaded into one channel of the AUC cell. To therefore make use of the interference system, NISTmAb and CsCl were loaded into the sample channel while the reference channel was loaded with the buffer used to prepare CsCl and in which the NISTmAb was dialyzed. The resulting interference signal (“fringes”) contains contributions from only the NISTmAb and CsCl.

A typical sample of NISTmAb and CsCl is shown in Fig. 2. Both absorbance (230 nm) and IF data are collected on the same AUC cell concurrently. The IF data are shown in Fig. 2A, while absorbance data are shown in Fig. 2B. Clearly, the interference dataset is not impacted by the peak at ∼6.8 cm that is visible in the absorbance dataset. This suggests that IF can be utilized to monitor the CsCl gradient, while absorbance optics can track the movement of the particle of interest. Scans taken at the same time points are displayed in Fig. 2A and Fig. 2B, and it is clear that the CsCl equilibrates more rapidly than NISTmAb. This can be quantified via a comparison of each scan with the final scan collected, as is shown in Fig. 2C. Monitoring of the gradient via interference optics is very insightful. As will be evident throughout this manuscript, this allows for characterization of the impact of various parameters on the shape of the gradient at equilibrium. Also of great importance is the visible isoconcentration point in Fig. 2A near 6.65 cm in the IF dataset. This would have been estimated historically but can now be determined experimentally using the IF optics.

**Figure 2.**
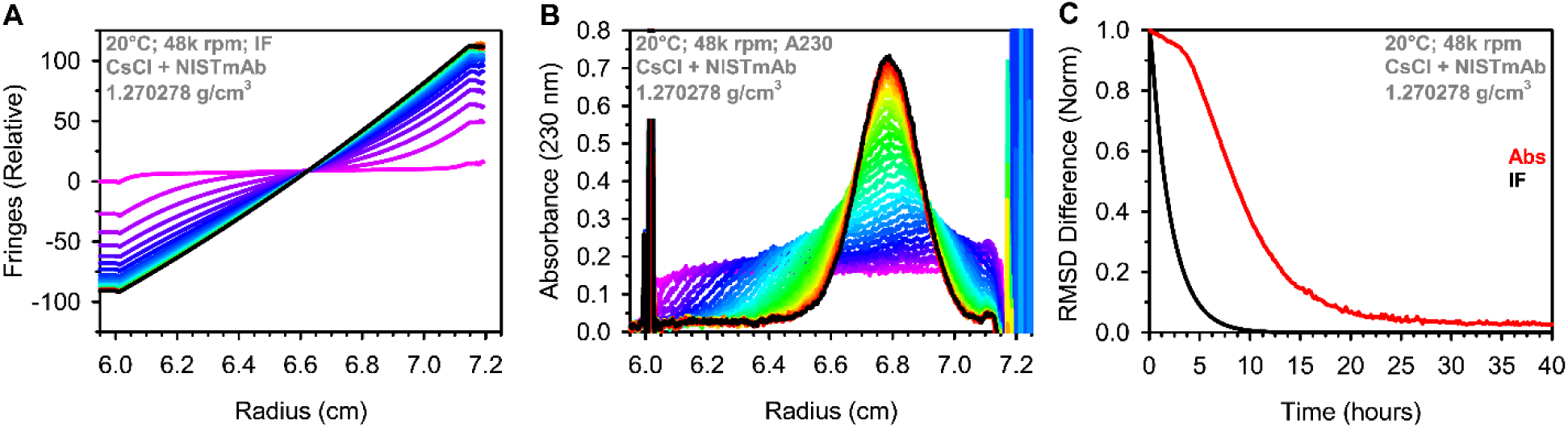
Formation of a CsCl density gradient in the AUC. (A) Interference scans collected over the first 24 hours of centrifugation. (B) Absorbance data (230 nm) collected over the first 24 hours of centrifugation. A total of 30 scans are shown in each dataset, where the rainbow color scheme reflects the time of the scan (magenta = first scan; red = last scan). The black line shows the final scan used for analysis. Each panel shows the relevant sample and instrument parameters. (C) The approach to equilibrium is shown as the RMSD difference compared to the final scan. Data were normalized to a maximum value of 1 to allow for easy visualization. The IF data shown in the black solid line reaches equilibrium much more quickly than the absorbance data shown in red.

### 3.3 Determination of NISTmAb buoyant density

#### 3.3.1 Effect of loading density

As already demonstrated in Figure 1, an analyte in a density gradient will equilibrate to a position in the cell where its buoyant density matches that of the solution density (i.e., the isopycnic point), yielding a peak within the AUC cell solution column. The focus of the presented method is to obtain multiple datasets wherein the particle forms a peak. Fig. 3 shows NISTmAb in six CsCl concentrations, each with slightly different solution densities as measured by a densitometer prior to loading the AUC cells. Both absorbance and interference data were collected. It is evident that the radial position of NISTmAb at equilibrium shifts with loading density (i.e., the solution density upon loading the AUC cell).

**Figure 3.**
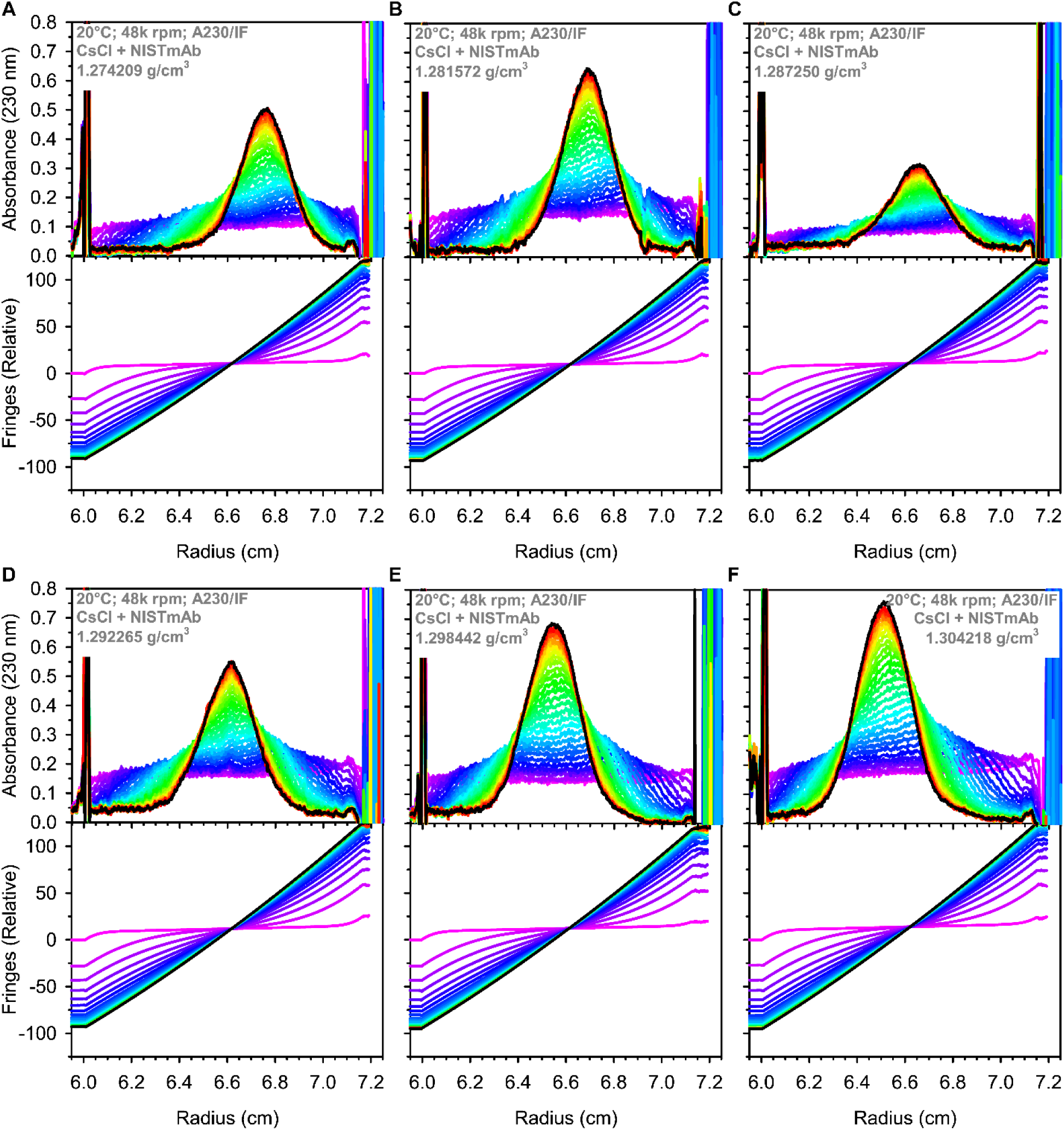
NISTmAb in CsCl density gradients at varying loading density. (A) – (C) show three lower CsCl loading densities, with absorbance data in the top plot and interference data in the bottom plot. Panels (D) – (F) show the three highest loading densities. All datasets were collected in a single experiment in an 8-hole rotor, along with the datasets in Figure 2A and Figure 2B. A total of 30 scans are shown in each dataset covering the first 24 hours of centrifugation, where the rainbow color scheme reflects the time of the scan (magenta = first scan; red = last scan). The black line shows the final scan used for analysis. Each panel shows the relevant sample and instrument parameters.

#### 3.3.2 Identification of the isoconcentration point

As stated previously, the primary goal of the presented method is to determine the buoyant density of a particle of interest without the use of a reference standard or model-dependent parameters. The critical component of this method is the identification of the isoconcentration point. The isoconcentration point is the radial position where the concentration of CsCl is the same at equilibrium as it was at the start of the experiment. The loading density of each sample (NISTmAb + CsCl) is directly measured using a densitometer, thus, the concentration and therefore density of the solution is known at the isoconcentration point. While Rossmanity and Mächtle focused on extrapolating the density across the entire cell via interference optics, the presented method uses <u>only</u> the density at the isoconcentration point. This avoids the difficulty in converting to absolute refractive index and accurately measuring the gradients near the sample meniscus and cell bottom [35]. This aspect helps ensure the model-independent nature of the presented method, because there are no assumptions made about the shape of the density gradient. In practice, the identification of the isoconcentration point is not trivial due to the inherent noises present in the interference detection system. The approach used to mitigate the inherent noise is described in Section “2.8 Interference data noise removal” and in Fig. S1. The IF datasets shown in Fig. 3 exhibit isoconcentration points near 6.65 cm, where the scans intersect. Estimation of the isoconcentration point has been used historically, however, this requires accurate and precise identification of the meniscus and bottom, and it includes assumptions about the gradient which the current method avoids [17,45]. A comparison of the interference-derived isoconcentration points and estimated ones is presented in Table S2.

#### 3.3.3 Buoyant density determination

With the ability to equate the measured loading density with the isoconcentration point at equilibrium in a DGE-AUC experiment, the question now focuses on obtaining a density for the NISTmAb peak positions. The radial position of NISTmAb and isoconcentration point was determined from each dataset shown in Fig. 2 and Fig. 3. These radii values are plotted in terms of the loading density of each sample (Fig. 4A). Firstly, one can readily observe that NISTmAb moves toward the meniscus (i.e., lower radius) as loading density increases. Secondly, the trend is essentially linear. This trend reflects the shape of the density gradient and will be explored further. Thirdly, there is some fluctuation in the isoconcentration point – this is an effect of having slightly different loading volumes in each AUC cell, which is unavoidable in practice.

**Figure 4.**
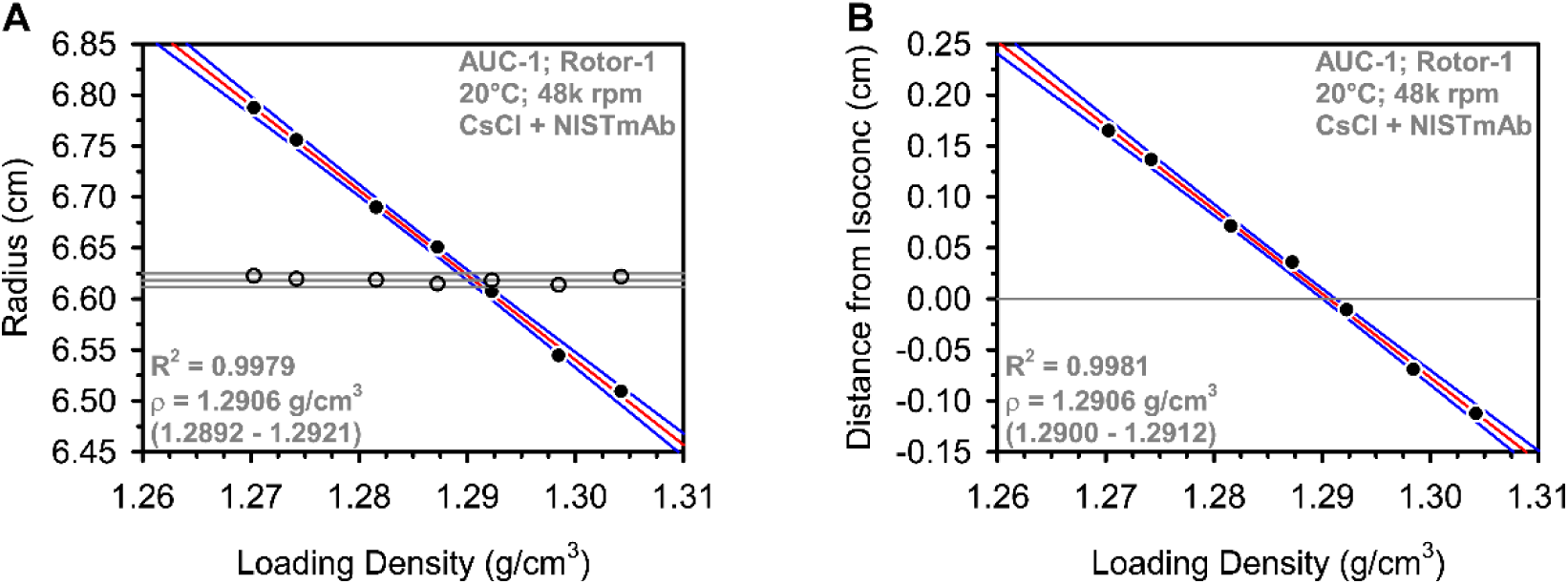
Buoyant density analysis of NISTmAb in CsCl at 48k rpm. (A) The datasets from Figure 2 and Figure 3 were analyzed to determine the radial position of the NISTmAb along the density gradient at each loading density. The corresponding radius and loading density are plotted in solid black markers, while the isoconcentration point from each interference (IF) dataset is shown as empty markers. The average isoconcentration point is shown as a solid grey line near 6.62 cm, with additional solid grey lines for ±2 standard deviations from the average. The data in (B) were obtained by calculating the difference between the observed NISTmAb radial position and the isoconcentration point of each sample shown in (A). The solid grey line represents the isoconcentration point. Linear regressions are shown as a red line, with the 95% confidence interval shown in blue lines. The fit statistics are shown in each panel, with the buoyant density 95% confidence interval in parenthesis. The 95% confidence interval in (A) also considered ±2 standard deviations along the average isoconcentration point radius.

Considering only the density at the isoconcentration point (i.e., the loading density) is taken with certainty in this method, the NISTmAb radii may be interpolated to find the loading density which would lead to the NISTmAb equilibrating exactly to the isoconcentration point. When this condition is satisfied, the NISTmAb buoyant density would be equal to the loading density. Based on the plot in Fig. 4A, this yields a buoyant density for NISTmAB in CsCl of 1.2906 g/cm^3^ with a 95% confidence interval of 1.2892 – 1.2921 g/cm^3^.

However, as mentioned, the isoconcentration point varies with sample volume loaded into the AUC cell. This incurs variability in the data which can be avoided if the data are plotted with respect to the isoconcentration point of each dataset, rather than plotting the absolute radii. To do this, the distance of the NISTmAb peak from the isoconcentration point is plotted for each sample. As seen in Fig. 4B, this results in an improved linear correlation coefficient. It also simplifies the interpolation, which can now be determined simply by performing a linear regression and reporting the x-intercept.

This yields a result of 1.2906 g/cm^3^ with a 95% confidence interval of 1.2900 – 1.2912 g/cm^3^. A value of 1.295 g/cm^3^ has been previously published for pooled human IgG in CsCl at pH 7.00 [46,47].

This demonstrates the basic approach of the model-independent method. Naturally, as with any global analysis, the number of datasets included and the approaches to error estimates can be adjusted as seen fit. It is also worth pointing out that there is no inherent assumption that the density gradient will be linear. In fact, a density gradient will always be evolving and changing with the concentration of the density gradient medium [17]. Any type of fitting (i.e., linear, polynomial, etc.) may be applied to the data presented in Fig. 4, however, with the instrumental parameters and the narrow window of densities chosen, the data are fitted very well to a linear regression. The IF data shown in Fig. 2 and Fig. 3 support the use of a linear regression, as the final scans themselves appear very linear.

### 3.4 Flexibility and adaptiveness of the presented method

To summarize this new method, a series of datasets are collected at different loading densities. The interference data are used to identify the isoconcentration point, while the absorbance data are used to monitor the peak position at equilibrium. Plotting the radial distance between the analyte position and isoconcentration point allows for simple and reliable determination of buoyant density (Fig. 4B). With a basic overview of the method now laid out, this section demonstrates how flexible the approach is in terms of instrumental and sample parameters. The results presented throughout this section also provide important insights into the nature of CsCl density gradients.

#### 3.4.1 Effect of rotor speed

Rotor speed has a major influence on the density gradient. This was noted early in the literature [17], which led to the convention of using standard conditions for DNA analysis [21,24]. To explore the use of the presented method across several rotor speeds, NISTmAb in CsCl was again examined across seven loading densities. The interference data for one sample at each speed was examined for its approach to equilibrium (Fig. 5A). Equilibrium of CsCl is reached within ∼10 hours at each speed, and after normalizing, there is no impact of rotor speed on the CsCl equilibration. When examining the interference scans at equilibrium, a significant impact on the slope of the gradient is observed (Fig. 5B). At higher rotor speeds, a steeper gradient is observed, meaning that a wider density range is captured within the solution column. The impact of this on NISTmAb is shown in Fig. 5C, where higher rotor speeds “compress” the peak into a narrower distribution. At lower rotor speeds, the peak is “spread out” over a larger range of the solution column. Because the peak shape and the density gradient change in a proportional way with respect to rotor speed, the resolving power between two given species is not expected to change [17], though we do not experimentally test this claim here. Additionally, rotor speed may need to be optimized based on the molecular weight of the particle of interest. For example, NISTmAb in CsCl at a rotor speed lower than 30,000 rpm may fail to produce a reliable distribution that can be analyzed. For the accurate determination of buoyant density using the presented method, it may prove advantageous to use a relatively low rotor speed to obtain a shallower density gradient, giving a smaller change in density per radial increment. On the contrary, if a mixture of particles is present, it may be advantageous to use a relatively high rotor speed to capture all particles within the solution column.

**Figure 5.**
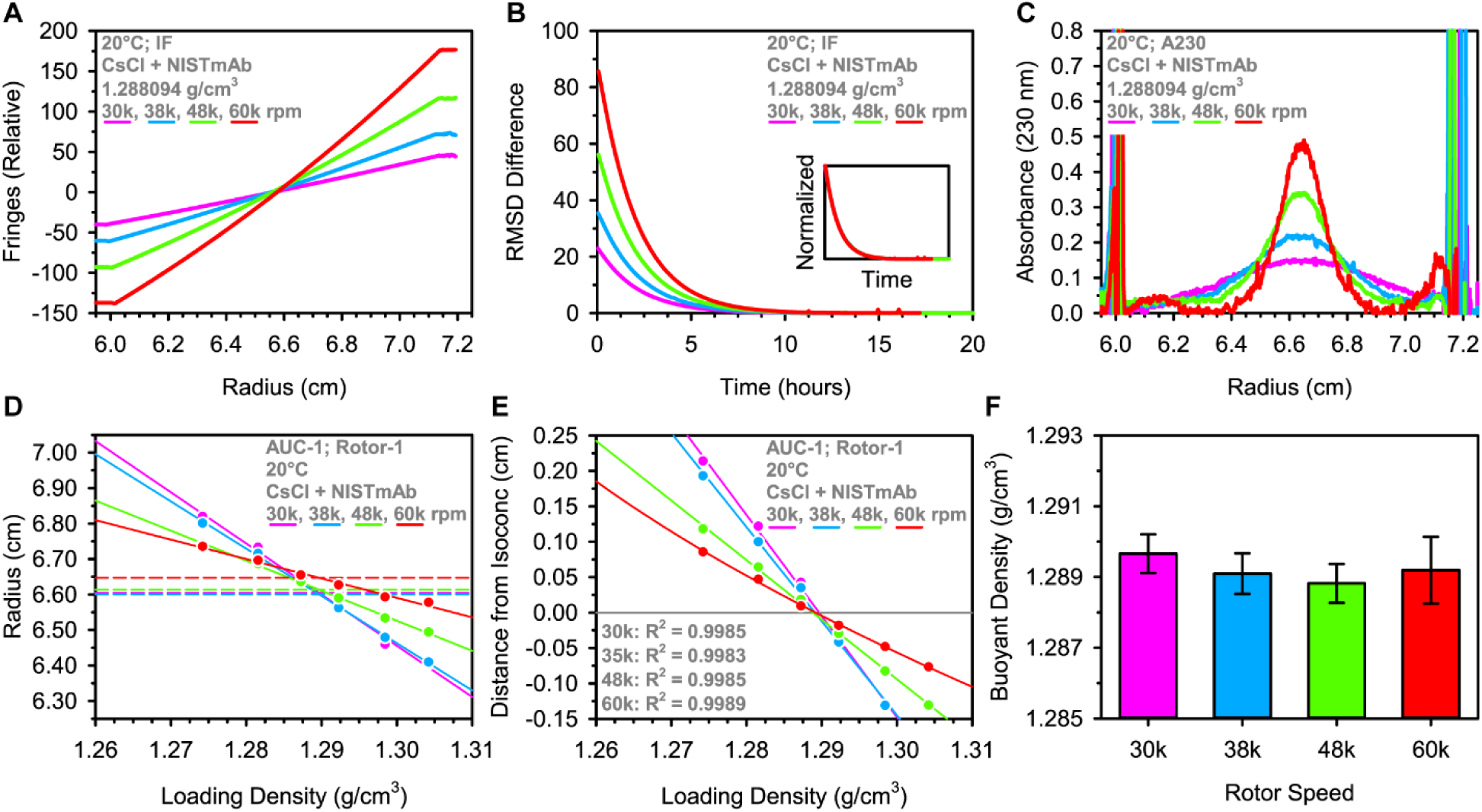
The impact of rotor speed on the analysis of NISTmAb in CsCl. The approach to equilibrium of CsCl at four rotor speeds is shown in (A), where the inset is normalized (all rotor speeds overlay). Scans taken at equilibrium are overlaid in (B). The corresponding absorbance data taken at equilibrium are shown in (C). Panels (A) – (C) contain data collected on the same sample at different rotor speeds (magenta = 30k, blue = 38k, green = 48k, red = 60k). Data shown for 60k rpm has been baseline corrected using a 3^rd^ order polynomial. (D) Shows results from NISTmAb in 7 loading densities at each speed. The solid lines are linear regressions, whereas the dashed lines are the average isoconcentration points. The distances from the isoconcentration points are shown in (E), with linear regressions and the corresponding correlation coefficients in the bottom left corner of the plot. The 60k rotor speed dataset in (E) was fitted to a 2^nd^ order polynomial instead, which resulted in an improvement in the R^2^. The resulting buoyant densities are summarized in the bar graph in (F), where error bars represent the 95% confidence interval.

Finally, radius vs loading density data were plotted for each rotor speed (Fig. 5D and Fig. 5E). At the highest rotor speed, a slight curvature is observed in the interference data (Fig. 5A; red). This is reflected in the slight curvature in the distance vs loading density plot (Fig. 5D; red). The 60k rpm dataset was fitted best with a 2^nd^ order polynomial regression, while all other data were highly linear and fitted with a linear regression. The resulting buoyant densities based on the distance from the isoconcentration point plot are very consistent and do not significantly change (Fig. 5E and Fig. 5F).

#### 3.4.2 Effect of cell misalignment

To demonstrate the robustness of this method in terms of providing reliable data for analysis, the AUC cells were intentionally misaligned in the rotor. The impact of this has been previously examined in the context of SV-AUC [48,49] and analytical band [9] techniques, where misalignment can cause significant errors in the results. Here, a set of seven samples was examined with and without intentionally misaligning the AUC cells. To allow for a more realistic test, the experiment with misaligned cells had each cell misaligned to differing degrees (randomly chosen between-5° and +5°). Based on visual inspection of the absorbance and interference data, cell misalignment causes an apparent shift in the isoconcentration point and NISTmAb peak position (Fig. 6A). This phenomenon is consistently observed across each misaligned cell (Fig. 6B).

**Figure 6.**
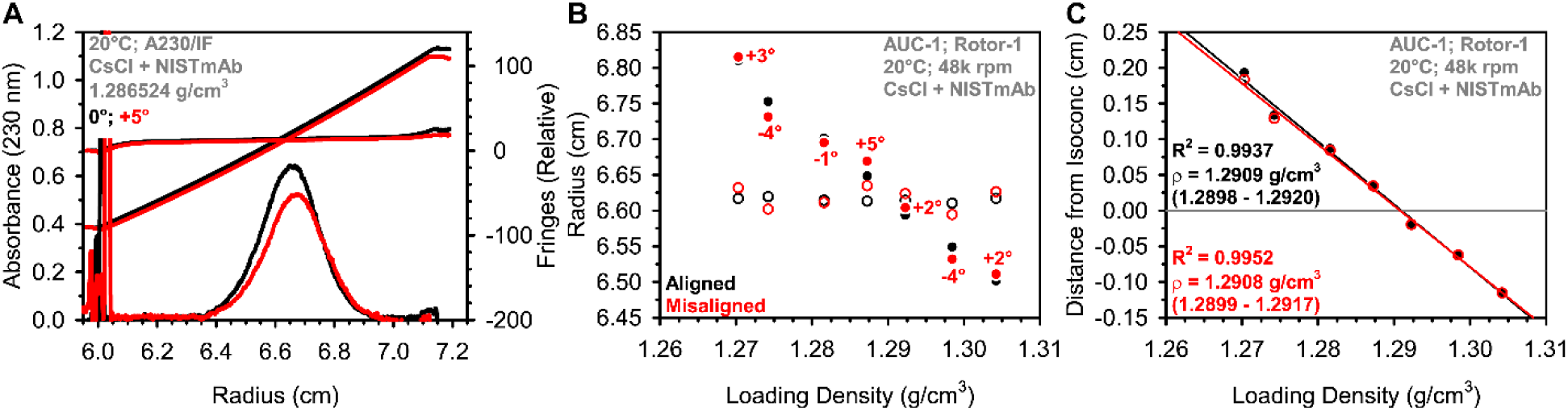
The impact of AUC cell alignment on the analysis of NISTmAb in CsCl. Panel (A) shows an overlay of absorbance scans taken of the same sample at equilibrium when the AUC cell was aligned (black) or misaligned by +5° (red). Also shown are the first and last interference scans (black = aligned, red = misaligned by +5°). The loading density vs radius results from NISTmAb in 7 loading densities are shown in (B) using the same color scheme (black = aligned, red = misaligned). Solid markers are NISTmAb radial positions, while empty markers are isoconcentration points. The distance from the isoconcentration point vs loading density results are shown in (C), with linear regressions shown as solid black and red lines, along with the corresponding correlation coefficients and buoyant densities (95% confidence intervals shown in parenthesis).

Remarkably, when measuring the distance from the isoconcentration point to the NISTmAb peak position, the effect on buoyant density is completely neglected (Fig. 6C).

#### 3.4.3 Effect of loading volume

The next parameter to examine is the sample loading volume. As mentioned previously, the exact isoconcentration point is influenced by the volume loaded into the AUC cell channels. The plotting of the distance from the isoconcentration point instead of the absolute radius appears to remove the consideration of loading volume. However, this was tested much more thoroughly in another DGE-AUC experiment summarized in Fig. 7. Seven samples of NISTmAb were again examined at different loading densities, but this time the volumes loaded into the AUC cells ranged from ∼210 µL to ∼420 µL. Examination of the interference data indicates that lower loading volumes result in the gradient reaching equilibrium faster, consistent with current knowledge about traditional sedimentation equilibrium experiments and previously reports for DGE-AUC [12]. The interference scans at equilibrium are shown in Fig. 7B, where the larger loading volume captures a wider range of fringes. This indicates that the density range of the gradient decreases as loading volume decreases, consistent with previous reports [15]. Importantly, when the scans are overlaid based on their isoconcentration points and relative intensity, the shape of the gradient is identical (Fig. 7B inset). This supports the use of the “distance from isoconcentration point” approach, because it shows that the density gradient is not significantly impacted by the loading volume.

**Figure 7.**
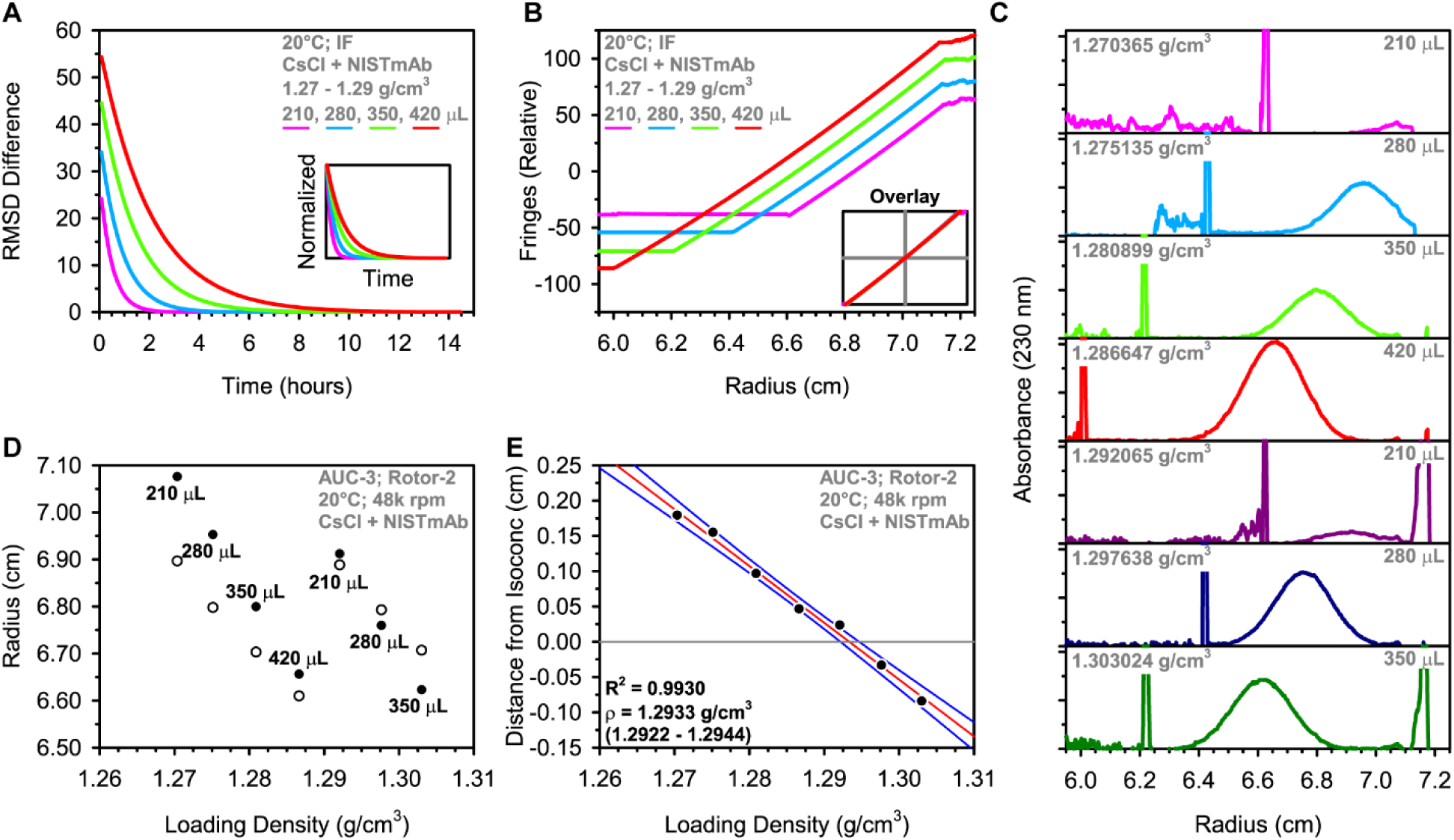
The impact of AUC cell loading volume on the analysis of NISTmAb in CsCl. The approach to equilibrium of CsCl at four loading volumes is shown in (A), where the inset is normalized (magenta = 210 µL, blue = 280 µL, green = 350 µL, red = 420 µL). Scans taken at equilibrium are shown in (B). The inset shows that the scans overlay very well when centered around the isoconcentration points indicated by the intersection of the grey lines. (C) Absorbance scans at equilibrium are shown in the stacked plot for all 7 samples of NISTmAb at different loading volumes – as indicated by labels in each plot. (D) The loading density vs radius results, where solid markers are NISTmAb radial positions and empty markers are isoconcentration points. The distance from the isoconcentration point vs loading density results are shown in (E), with the linear regression shown as a red line and with blue lines showing the 95% confidence interval. The correlation coefficient and buoyant density (95% confidence interval shown in parenthesis) are indicated by the text within the plot.

The absorbance data are shown in Fig. 7C and provide several important insights. Firstly, the area under the distribution and corresponding peak amplitude decreases greatly with loading volume, despite each sample having a similar NISTmAb concentration. This can be explained by the concentrative nature of the DGE-AUC experiment – the total quantity of NISTmAb in each cell is the relevant parameter to the peak signal area. The particle of interest is concentrated within the density gradient, forming a peak. This illuminates the use of preparative density gradient ultracentrifugation that is popular for virus purification [28,29]. Secondly, the absolute radial position of NISTmAb is useless in the radius vs loading density plot (Fig. 7D).

Examining the distance from the isoconcentration point approach once again proves powerful in recovering a buoyant density (Fig. 7E) that is consistent with that determined on the same instrument and rotor using similar loading volumes (Fig. S2).

### 3.5 Application to DNA

Historically, DNA has possibly been the most popular application of DGE-AUC. To demonstrate the use of the presented method for a particle other than NISTmAb, experiments were conducted on λ bacteriophage DNA in CsCl. An example of the λ DNA absorbance data is shown in Fig. 8A. The peak is extraordinarily sharp, owing to the high molecular weight of λ DNA (∼30 MDa), meaning that this approach also requires a very small sample quantity. Two separate DGE-AUC experiments were performed with λ DNA, one at standard conditions (44,000 rpm and 25°C) and one at 48,000 rpm and 20°C. Also, these were performed without a reference standard that is typically required [21,27]. The results indicate good linear regression fits and tight confidence intervals around ∼1.7 g/cm^3^, consistent with ranges of buoyant density for DNA in CsCl (Fig. 8B and Fig. C) [24]. There is a slight difference between the buoyant density under each condition, but this can be attributed to temperature-dependent changes in the hydration of DNA [50].

**Figure 8.**
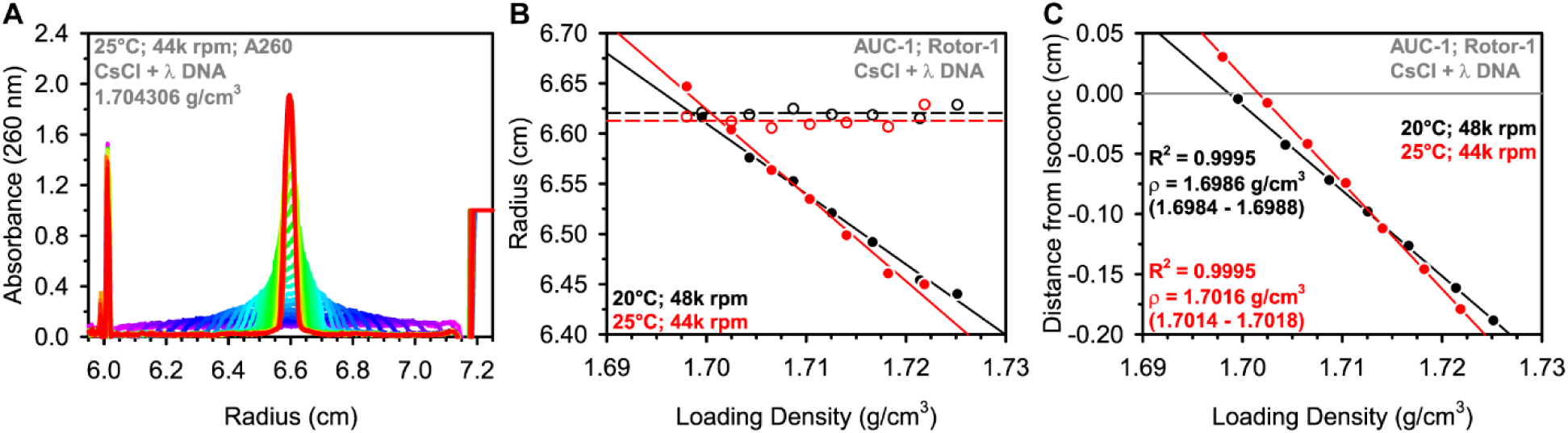
The analysis of λ DNA in CsCl. (A) Absorbance data (260 nm) showing the equilibration of λ DNA in CsCl. A total of 30 scans are shown, where the rainbow color scheme reflects the time of the scan (magenta = first scan; red = last scan). (B) Loading density vs radius results, where solid markers are λ DNA radial positions and empty markers are the isoconcentration points (black = 20°C, 48k rpm; red = 25°C, 44k rpm). Linear regressions are shown in solid lines. (C) The distance from the isoconcentration point vs loading density results for the two datasets are shown. Linear regressions are shown in solid lines, with the correlation coefficients and buoyant densities (95% confidence intervals shown in parenthesis) are indicated by the text within the plot.

### 3.6 Application to PS beads in sucrose

To further demonstrate the flexibility of the presented method, PS beads were also examined in sucrose. Here, we prepared samples in sucrose rather than CsCl, because the ionic nature of CsCl causes agglomeration of the PS beads. Additionally, these data were collected on the newer Optima AUC instead of the XL-I and used 3 mm titanium centerpieces instead 12 mm aluminum centerpieces. PS is a well characterized material with a known density of ∼1.055 g/cm^3^ [51,52]. Data collected using the absorbance and interference detection systems are shown in Fig. 9A – Fig. 9C. Once again, a very sharp peak is observed, consistent with the high molecular weight of the beads. Due to the high refractive index of PS, the interference scans show a distinct peak from the PS beads. This opened the possibility of using only interference optics for the method, rather than requiring that the particle of interest has a chromophore which absorbs light at a given wavelength. In fact, the signal tracked in this experiment was caused by light scattering rather than absorption.

**Figure 9.**
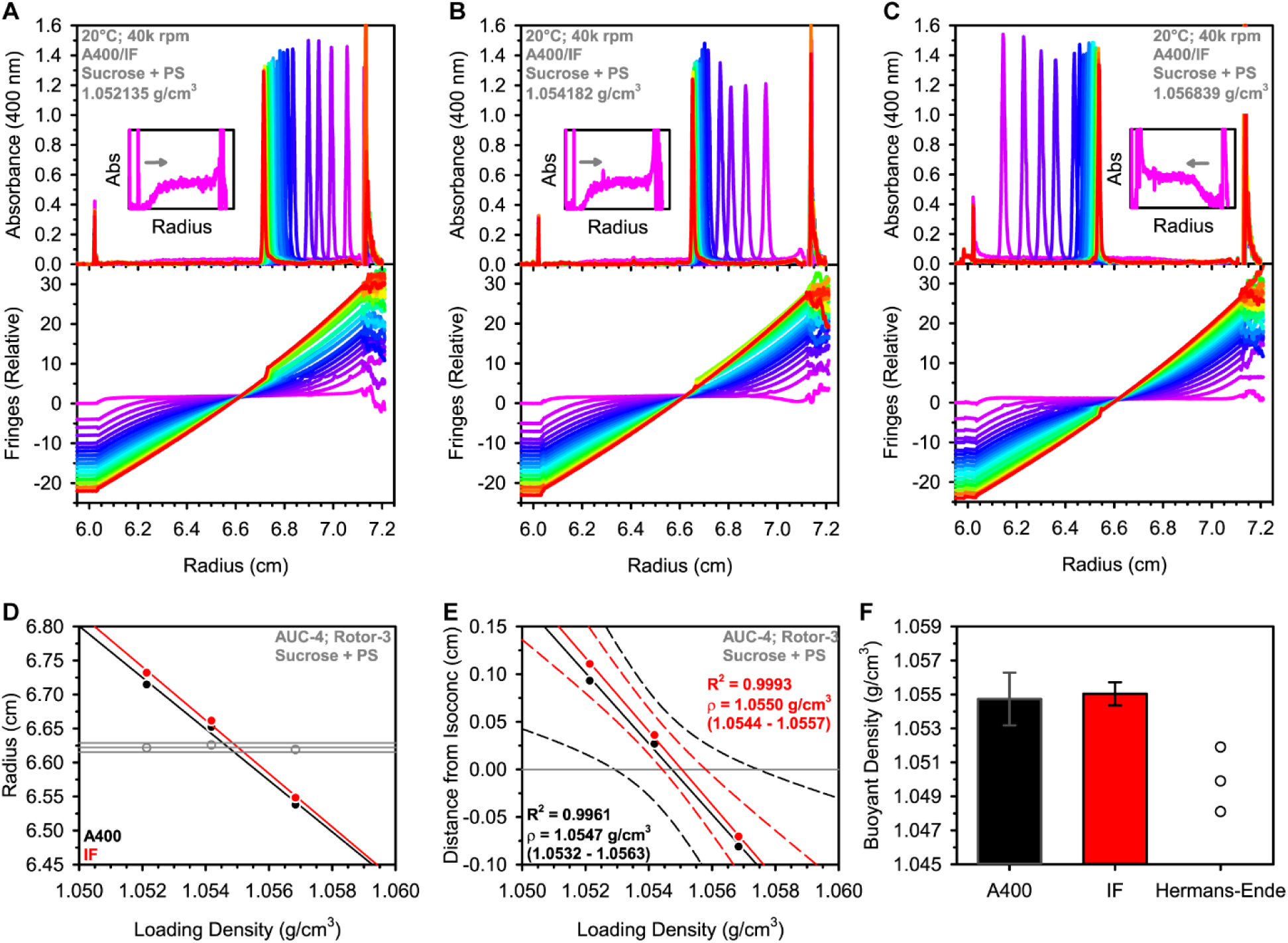
Polystyrene beads in sucrose density gradients at varying loading density. (A) – (C) show three loading densities, with absorbance data in the top plot and interference data in the bottom plot. A total of 30 scans are shown in each dataset covering the first 24 hours of centrifugation, where the rainbow color scheme reflects the time of the scan (magenta = first scan; red = last scan). The insets show a zoom of the first scan, showing either sedimentation or floatation at the start of the experiment (see direction of grey arrow). (D) The loading density vs radius results from the three datasets, where solid markers are PS radial positions and empty markers are isoconcentration points. The black markers correspond to analysis of the absorbance data, while the red markers used only the interference data. The distance from the isoconcentration point vs loading density results are shown in (E), with the linear regressions shown as solid lines and dashed lines showing the 95% confidence interval. The correlation coefficient and buoyant density (95% confidence interval shown in parenthesis) are indicated by the text within the plot (black = absorbance data, red = interference data). (F) A bar graph summarizing the buoyant density results determined via the presented method using absorbance data or interference data for determining the radial position of PS (error bars show the 95% confidence interval), or the classical Hermans-Ende method using absorbance data (individual results from local analysis). This method provides an independent result from each of the three datasets.

The results were examined based on the previously performed approach (absorbance for PS peak determination and IF for isoconcentration determination), as well as using only the interference data. As can be seen, the results are consistent from each approach. The interference data alone resulted in slightly better fit statistics, but both buoyant density values are in very good agreement with the literature value. This experiment demonstrates the effectiveness of the method for synthetic particles and for other density gradient media. Importantly, it also confirms that this approach can be extremely accurate.

As a comparison to a traditional method, the Hermans-Ende method [33] was applied to the same datasets. This approach is a local analysis (i.e., one dataset is analyzed at a time). The details of this method are discussed in the section: “2.9 Hermans-Ende analysis”. The results from the analysis of each dataset are plotted in Fig. 9F. Individual estimates of the buoyant density from the Hermans-Ende method ranged from 1.0481 – 1.0519 g/cm^3^ (Table S1), each of which are significantly less than that determined by the present method.

Closer inspection of the raw data reveals significant information on the buoyant density. The insets in Fig. 9A – Fig. 9C show the first absorbance scan obtained. Sedimentation is readily observed for the two lower loading densities, but floatation (migration toward the meniscus) is observed for the higher loading density. Considering that the experiment starts with a constant density across the cell (i.e., the loading density), this observation indicates that the buoyant density of the PS beads must be greater than 1.0542 g/cm^3^ but less than 1.0568 g/cm^3^.

A notable observation for the PS beads is the formation of a sharp peak which then equilibrates to the final radial position. In contrast, the NISTmAb and DNA both move to the final radial position while the peak forms. This may be explained by the extraordinary size of the PS beads and their resulting low diffusion coefficient. These two characteristics mean that the particles sediment or float very rapidly (at this rotor speed), concentrating them to the cell bottom or meniscus. Then, as the sucrose density gradient begins to form, the PS beads migrate to their respective isopycnic position within the cell.

### 3.7 In-depth examination of the presented method

#### 3.7.1 Instrument-to-instrument variability

A multi-laboratory SV-AUC comparison has previously been performed using bovine serum albumin [53]. This provided useful insights into sources of variability between SV-AUC measurements. To start to examine variability in DGE-AUC, a total of three Beckman Coulter XL-I AUCs and two 8-hole rotors were utilized for NISTmAb in CsCl at 48,000 rpm and at 20°C. The results from these experiments are shown in Fig. S2 and show excellent agreement between two of the AUCs with different rotors. The third AUC showed a shift in the buoyant density beyond the 95% confidence interval of the other results. This could be attributed to a discrepancy in the radial calibrations of the absorbance and interference systems. A radial correction was applied to the absorbance and interference data by shifting the scans left or right to align the counterbalance edges observed during this experiment and the experiment on AUC-1. Essentially, this makes use of the counterbalance as an internal standard for radial position, and the results after radial correction of AUC-3 data were consistent with the other instruments. Alternatively, some particles might be more readily analyzed using only interference data, as was shown in Fig. 9 for PS. This would avoid any concerns about the agreement between the two optical systems.

#### 3.7.2 Wiener skewing and the accuracy of radial measurements in a density gradient

The refraction of light is dependent on the density of the material encountered. In the context of density gradient experiments, the light from the AUC optical source will not encounter a consistent density across the cell. In theory, there will be differential refraction at every point along the density gradient in the AUC cell. This will result in light refracting more and more as it encounters higher density solution. The result of this is that the light reaching the detector has been shifted to a higher radius. This has been explored previously in the context of SV-AUC and sedimentation equilibrium AUC experiments, and the effect was determined to be negligible unless an obvious “dark band” is observed – a region where no light reaches the detector [54].

Theoretically, factors impacting Wiener skewing are the optical system’s focal plane within the AUC cell and the magnitude of the refractive index gradient [54]. The focus of the optics is not readily adjustable by the user. The magnitude of the refractive index gradient may be mitigated in part by using 3 mm pathlength centerpieces rather than the standard 12 mm pathlength centerpieces [13]. The effect is also reduced at lower speeds, because the density gradient becomes shallower.

Experimentally, the degree of Wiener skewing can be measured very simply in the context of a particular experimental setup. In Fig. S3, the degree of Wiener skewing observed was evaluated by drawing lines on the top AUC cell window using a black ballpoint ink pen. The lines provide reference points, not unlike the more sophisticated approaches used by others [54,55]. The positions of the lines were monitored by absorbance and interference optics at a rotor speed of 60,000 rpm – where the effect should be most significant. At the beginning of the experiment, there is essentially no density gradient and therefore the accurate radial position of the lines should be obtained. However, as the density gradient forms, the radial position of the lines may shift due to Wiener skewing. The radial position of the lines was monitored for any shift throughout the experiment by absorbance and interference optics. Similar, minor shifts (<0.01 cm) were observed in both detection systems at 60,000 rpm and in 12 mm centerpieces – where the effect should be most extreme.

Interestingly, because the isoconcentration point is the fundamental reference point for the DGE-AUC method, Wiener skewing should have little to no impact on the resulting buoyant density so long as the two optical systems are impacted similarly (or in the case where only interference is being used). In conclusion, any major Wiener skewing is likely to have an insignificant impact on buoyant density determination due to the nature of the presented DGE-AUC method – the comparison of relative positions with respect to the *observed* isoconcentration point. Nonetheless, its presence may be readily examined for any given experimental setup.

#### 3.7.3 Impact of the particle on solution density

Measurement of the loading density is clearly very important to this method. In the current work, the density was measured directly using a densitometer. Others have used refractometers to measure the refractive index of CsCl or other gradient forming media, which is then converted to a concentration and then to a density [13,26].

Refractometry has the advantage of being quick and using very small sample volumes; however, it assumes that the density is dictated only by the gradient forming material.

An interesting aspect of the density measurement of the sample is that while the particle of interest can contribute to the bulk density of the sample, this impact would be minimized as the solution density nears the particle’s buoyant density. Therefore, the presented method of interpolation should not be negatively impacted by cases where the particle has a significant impact on the bulk density of the sample.

#### 3.7.4 Considering pressure effects

The analytical ultracentrifuge is capable of yielding pressures inside the sample cell of at least up to ∼350 bar [56]. There are several ways that pressure may impact DGE-AUC experiments [19,46,57]. (1) The solution itself may become compressed into a smaller volume, such that the loading density as measured in a density meter becomes an underestimate of the density under the conditions of the first scan. This would lead to an overestimate of the buoyant density using the presented method, with the effect being larger as rotor speed is increased. The extent of this effect may be evaluated directly in the AUC for a given solution using a relatively simple approach for determining solution compressibility – measure the change in volume of the solution across different rotor speeds [56]. (2) There may exist an impact on the dispersion of gradient forming material across the solution column due to pressure. This may cause a significant effect on the resulting buoyant density when using approaches that predict the density distributions based on methods of Ifft and colleagues [17,57]. In the presented method, this pressure effect is not a concern due to the measuring of the density gradient directly via the interference optics. (3) Pressure may impact the structure or solvation of the particle of interest, modulating the buoyant density [19,46,47,57]. Pooled human IgG showed no significant change in buoyant density with respect to pressure, while bovine serum albumin exhibited an increased buoyant density of ∼0.005 g/cm^3^ per 100 atm [47]. The current work on NISTmAb, a recombinant humanized IgG1κ, is consistent with the human IgG data – no significant impact of pressure on buoyant density as evident by the rotor speed data presented in Fig. 5F.

#### 3.7.5 Application to molecular weight determination

The apparent molecular weight of a given particle in a density gradient may be estimated [16,44,58]. Traditionally, this requires an accurate value for the density gradient, (*dρ/dr*)_0_, at the radial position of the particle at equilibrium, along with the particle’s buoyant density. The density gradient parameter has traditionally been determined by a constant (*β*) that is intrinsic to the density gradient medium, and depends on angular velocity and radial position [44]. These values are tabulated for given conditions [17,45,57,59]. Interestingly, the presented method determines the density gradient parameter via the slope of the radius vs loading density plots. These directly report the change in density per the change in radius, given that the buoyant density of the particle remains unchanged. Therefore, the determined slope may be used in the equation for determining apparent molecular weight (*M*_app_). The *M*_app_ value for NISTmAb in CsCl was estimated (see section “2.10 Molecular weight determination” for more details) from data collected during the previously discussed experiments (Fig. 10A). The resulting *M*_app_ was 148.1 kDa (144.6 – 151.7). Each relevant parameter was investigated for its potential to propagate error to the result. The density gradient parameter caused the greatest uncertainty of approximately ±4 kDa. It should be noted that this molecular weight is an “apparent” molecular weight that includes the mass of NISTmAb (∼145 kDa) and any associated Cs^+^ ions, the latter of which also impact the buoyant density of the particle compared to the particle density in water.

**Figure 10.**
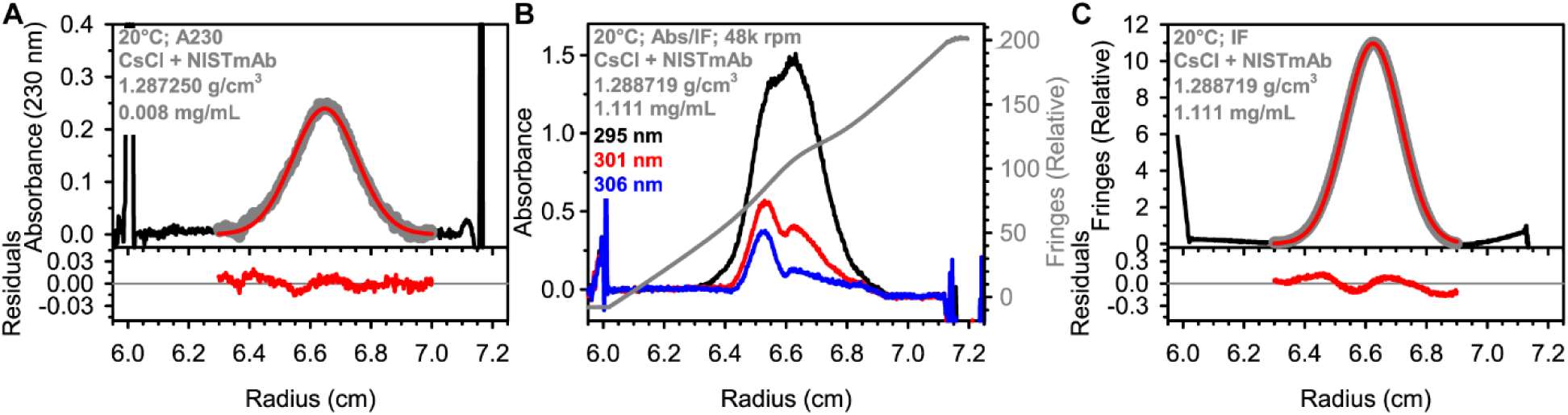
Molecular weight analysis of NISTmAb in CsCl. (A) Fitting of NISTmAb at a low concentration (0.008 mg/mL) via absorbance data (230 nm) and the equation described in the section “2.7 Molecular weight determination”. The black solid line in the upper panel is a scan taken at equilibrium and adjusted via a linear baseline correction. The grey symbols represent the range of data fitted to the Gaussian equation, with the red line being the best fit. The lower panel shows the residuals from the fit. Panel (B) shows absorbance scans collected at several wavelengths for a sample at 1.111 mg/mL. The interference scan is shown as a grey line and on the right y-axis. Panel (C) shows fitting of the IF data from (B) after a polynomial baseline correction to determine M_app_ [197.1 kDa (192.5 – 201.9)].

#### 3.7.6 Exploring concentration-dependence

An additional consideration is that the shape of the particle’s distribution may be impacted by concentration-dependent thermodynamic nonideality [60]. One may also be interested in concentration-dependent interactions or associations. To demonstrate this potential application, NISTmAb was characterized by DGE-AUC at >1 mg/mL – an approximately 100-fold increase over previous samples. Data were collected at several absorbance wavelengths (Fig. 10B). Because the sample reaches equilibrium in the AUC, any number of wavelengths may be scanned to obtain a distribution that is within the linear range of the absorbance optics. Interestingly, the absorbance scans showed major abnormalities in the peak shape of the absorbance scans that are likely due to Wiener skewing. The interference scan does, however, present a reasonably shaped peak, which could be evaluated for molecular weight determination (Fig. 10C). An increase in apparent molecular weight was observed at this concentration.

Thermodynamic nonideality is expected to cause an underestimate of the molecular weight [60]; however, there may exist some self-association of NISTmAb at such high concentrations, even in the presence of CsCl, that could compete with nonideality.

## 4. Conclusions

Density gradient equilibrium AUC (DGE-AUC) has a rich history and has many modern applications. The method presented here demonstrates a fundamental approach to determining the buoyant density of a particle of interest using either absorbance and interference detection systems, or in the case of PS beads, interference only. The method is model-independent, highly accurate, precise, robust, and does not require a standard reference material or calibration curve. Importantly, it can be readily applied to many different systems, especially those with heterogeneous composition. While not explored in the current work, this approach is not limited to homogeneous systems – multiple species may be analyzed concurrently. It should be noted that DGE-AUC-derived buoyant density includes hydration/solvation effects, and the density gradient medium may interact with the particle of interest.

The importance of the model-independent nature of the presented method cannot be overstated. Traditional methods for analyzing particles by DGE-AUC required a model of the density gradient. They often involved simplifying assumptions, such as ideality of the gradient forming medium, and running at standard conditions and with reference markers. The analyses were performed on single datasets, rather than multiple datasets as with the presented method. Global analyses provide additional rigor and allow the calculation of error estimates.

There has long been interest in simulating density gradient and macromolecular band formation [13,61–63]. The limiting factor of such approaches is likely to be the shape of the density gradient, including the accurate prediction of the isoconcentration point [45,57]. In the presented method, the isoconcentration point is directly measured using the interference detection system found on modern AUCs. This approach may be invaluable to informing future simulations and model-dependent analyses.

## Abbreviations

AAV: Adeno-associated virus
mAb: Monoclonal antibody
AUC: Analytical ultracentrifugation
CsCl: Cesium chloride
DGE: Buoyant density gradient equilibrium λ Lambda bacteriophage
PS: Polystyrene
SDS: Sodium dodecyl sulfate
SV-AUC: Sedimentation velocity analytical ultracentrifugation

## Acknowledgements

The authors would like to acknowledge Dr. John W. Burgner 2^nd^ for useful discussions in the early stages of this work. PCL and JW would like to acknowledge Gudrun Bleyer and the Vogel lab for providing the PS beads as well as Sandra Wittpahl, Hadi Soltanmoradi and Tzu-Ling Su for their support in experiments.

## Data Availability

Data series are supplied in a supplementary file.

## CRediT authorship contribution statement

AEY

- Formal analysis

- Methodology

- Data curation

- Visualization

- Writing – original draft

PCL

- Data curation

- Formal analysis

- Writing – original draft

JW

- Resources

- Writing – review and editing

MTD

- Methodology

- Writing – review and editing

LNP

- Methodology

- Resources

- Writing – review and editing

