## Supplemental_Materials for "A Fundamental Approach to Buoyant Density Determination by DGE-AUC"

### Supplementary material

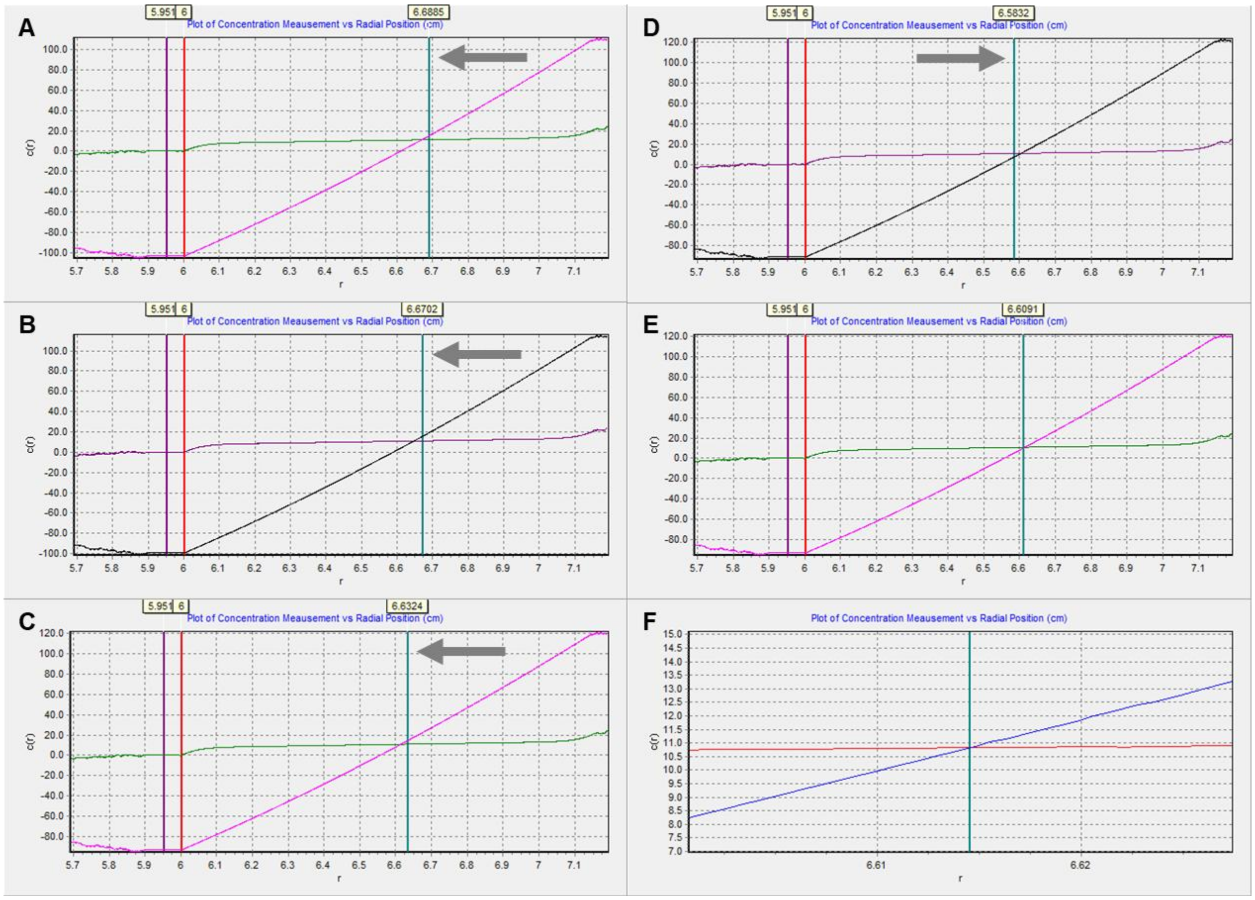

**Figure S1. Robustness of the identification of the isoconcentration point within SEDVIEW.** Screenshots of SEDVIEW are presented for the identification of the isoconcentration point of a typical interference dataset from NISTmAb in CsCl at 48,000 rpm. The first scan and a scan at equilibrium are shown. The solid vertical purple line is the radius chosen for the jitter adjust. The vertical red line is the meniscus radius, which is irrelevant for the identification of the isoconcentration point and is ignored here. The vertical teal line is the radius chosen for the integral fringe removal. Panels (A) – (D) show radii chosen and the subsequent scan positioning. For (A) – (C), the radius chosen resulted in an intersection of the scans to the left (i.e., lower radius) of the teal vertical line. In each iteration, the teal line was moved to a lower radius, and the intersection kept moving to a lower radius. Eventually, as shown in (D), the intersection moved to a higher radius than the teal line. The teal line was then moved back toward the intersection in finer increments until the vertical teal line was perfectly located at the intersection (E). Panel (F) shows the zoom function in SEDVIEW, which allows for extremely precise identification of the isoconcentration point as the intersection of the first and final scans.

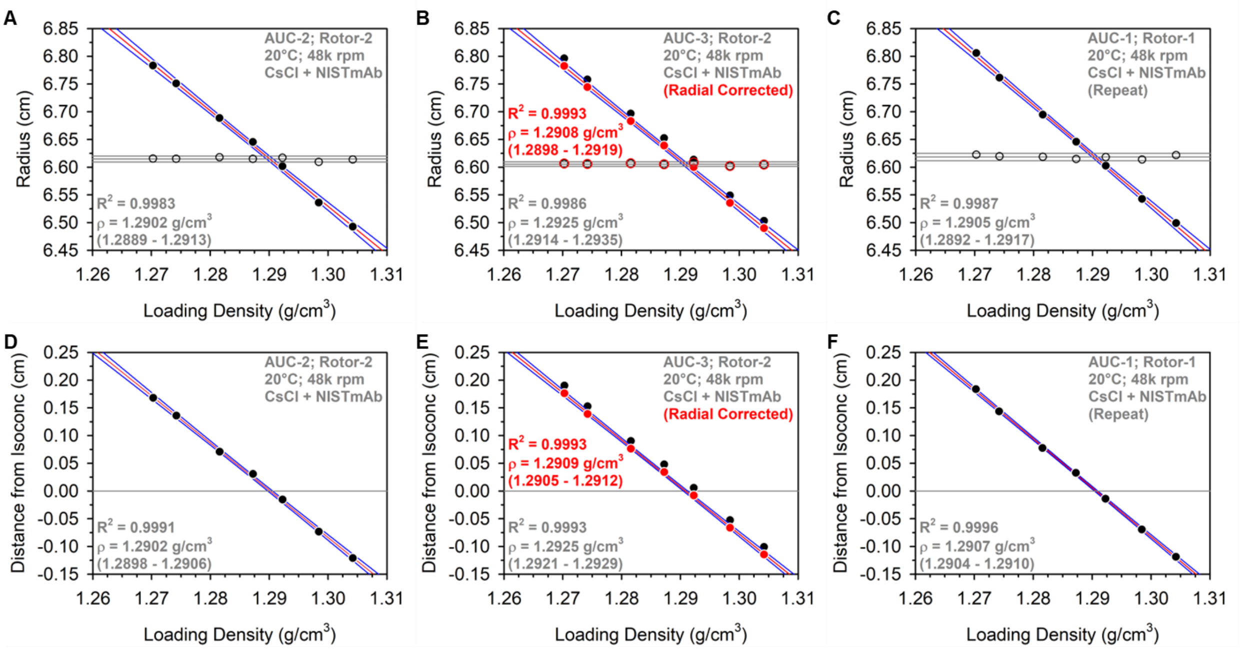

**Figure S2. Buoyant Density analysis of NISTmAb in CsCl at 48k rpm on different instruments.** The samples from Figure 4 were mixed and re-analyzed on two other instruments (AUC-2 and AUC-3) and one other 8-hole rotor (Rotor-2). The radial position (filled black marker) is plotted for each loading density in (A) and (B) for the two additional experiments, along with the isoconcentration point (empty black marker; average shown as grey line with additional solid grey lines for ±2 standard deviations from the average). (C) Afterwards, the samples were re-analyzed on AUC-1 with Rotor-1 ("Repeat") to ensure no changes in the samples occurred over the course of testing across instruments. Panels (D) – (F) show the difference between the observed NISTmAb radial position and the isoconcentration point of each sample. The solid grey line represents the isoconcentration point. Linear regressions are shown as a red line, with the 95% confidence interval shown in blue lines. The fit statistics are shown in each panel, with the buoyant density 95% confidence interval in parenthesis. In panels (B) and (E), the red data and text display the results after correcting for inconsistencies in the radial calibration between the absorbance and interference systems on AUC-3. For clarity, the linear regression and 95% confidence interval are shown only for the corrected data.

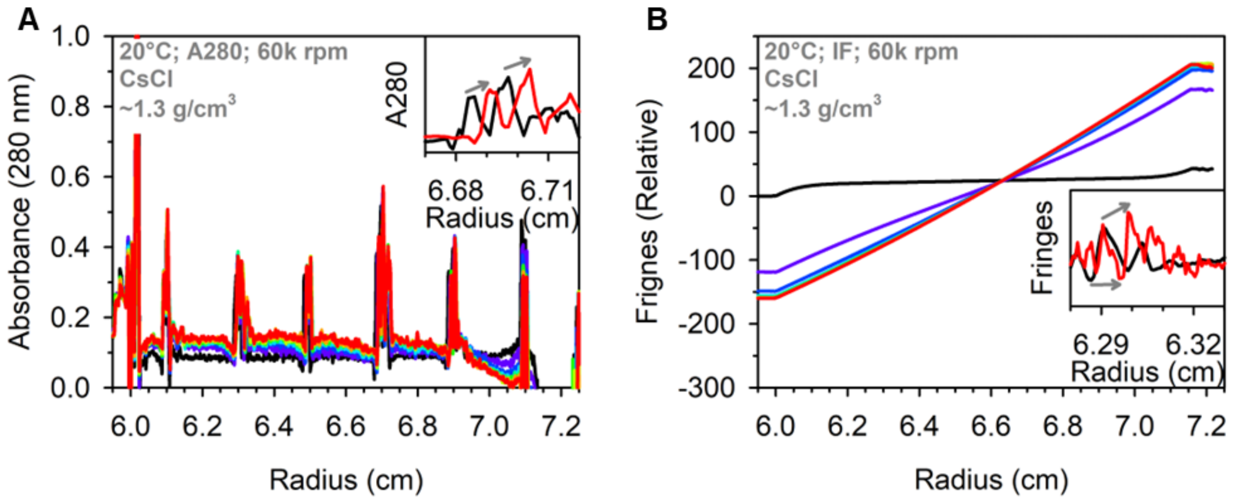

**Figure S3. Impact of refractive index effects in a CsCl density gradient.** A solution of CsCl was loaded into a cell assembled with windows which had been marked on the bottom window to yield noise in the scans at consistent radial positions. Panel (A) shows absorbance scans taken over the course of CsCl equilibration (first scan = black, last scan = red). Corresponding interference scans are shown in (B). The insets highlight a specific region of the cells where distinct artifacts can be identified in the first scan (black line) and at equilibrium (red line). Grey arrows illustrate minor shifts (~0.006 cm) in the radial positions that may be attributable to refractive index effects known as Wiener skewing. Polynomial baseline corrections were applied to the interference scans to obtain a useful overlay of the data. Scans were collected at 60,000 rpm, where the density gradient, and therefore Wiener skewing effect, is expected to have the most dramatic effects.

**Table S1. Experimental parameters used for Hermans-Ende analysis of PS beads.**

| | | | |
|---|---|---|---|
| **Temperature (K)** | 293 | 293 | 293 |
| **Angular velocity ($s^{-1}$)** | 666.6667 | 666.6667 | 666.6667 |
| **Loading density ($g/cm^3$)** | 1.0521 | 1.0542 | 1.0568 |
| **Meniscus (cm)[a]** | 6.0210 | 6.0219 | 6.0225 |
| **Bottom (cm)[b]** | 7.1097 | 7.1134 | 7.1050 |
| **Peak position (cm)** | 6.7138 | 6.6518 | 6.5379 |
| **Loading volume (μL)** | 110 | 110 | 110 |
| **Density of low-density material ($g/cm^3$)** | 0.9982 | 0.9982 | 0.9982 |
| **Compressibility of low-density material ($Pa^{-1}$)** | $5\times10^{-10}$ | $5\times10^{-10}$ | $5\times10^{-10}$ |
| **Density of gradient forming material ($g/cm^3$)** | 1.5518 | 1.5518 | 1.5518 |
| **Compressibility of gradient forming material ($Pa^{-1}$)** | 0 | 0 | 0 |
| **Molar mass of gradient forming material (kg/mol)** | 0.3423 | 0.3423 | 0.3423 |
| **Buoyant Density ($g/cm^3$)** | 1.0481 | 1.0499 | 1.0519 |

[a]The meniscus position was taken as the peak of the positive optical artifact in the absorbance dataset.
[b]The bottom position was taken as the point where the light intensity decreases in the absorbance dataset.

**Table S2. A comparison of the identification of the isoconcentration point.**

| | | | | | | | | |
|---|---|---|---|---|---|---|---|---|
| **Loading Density (g/cm$^3$)** | 1.270278 | 1.274209 | 1.281572 | 1.28725 | 1.292265 | 1.298442 | 1.304218 | **Buoyant Density (g/cm$^3$)** |
| **Meniscus (cm) [Abs]**[a] | 6.0182 | 6.0152 | 6.0142 | 6.009 | 6.0155 | 6.0048 | 6.0154 | |
| **IF-determined Isoconcentration** | 6.6225 | 6.6195 | 6.6185 | 6.6146 | 6.6179 | 6.6137 | 6.6216 | 1.2906 (1.2900 – 1.2912) |
| **Bottom (cm) [geometric]** | 7.15 | 7.15 | 7.15 | 7.15 | 7.15 | 7.15 | 7.15 | |
| **Calculated Isoconcentration**[b] | 6.6084 | 6.6070 | 6.6066 | 6.6042 | 6.6071 | 6.6023 | 6.6071 | 1.2921 (1.2915 – 1.2927) |
| **%Change Isoconcentration**[c] | 0.2133% | 0.1887% | 0.1805% | 0.1574% | 0.1625% | 0.1727% | 0.2190% | |
| **Bottom (cm) [rotor stretch]**[d] | 7.16896 | 7.16896 | 7.16896 | 7.16896 | 7.16896 | 7.16896 | 7.16896 | |
| **Calculated Isoconcentration** | 6.6186 | 6.6173 | 6.6168 | 6.6145 | 6.6174 | 6.6125 | 6.6174 | 1.2909 (1.2903 – 1.2914) |
| **%Change Isoconcentration**[c] | 0.0583% | 0.0336% | 0.0254% | 0.0022% | 0.0074% | 0.0174% | 0.0640% | |

[a]The meniscus position was taken as the peak of the positive optical artifact in the absorbance dataset.

[b]The isoconcentration point was calculated using equation 12a from Ifft et al. 1961 [1]: $\sqrt{(r_b{}^2 + r_a{}^2)/2}$ where $r_b$ is the bottom position and $r_a$ is the meniscus position.

[c]The percent change in the isoconcentration point was calculated as: (|Calc Isoconc – IF-determined Isoconc|/IF-determined Isoconc) × 100.

[d]The bottom position was corrected for speed-dependent rotor stretch in UltraScan [2]. Rotor stretch coefficient 1 was 1.18423e-07 and rotor stretch coefficient 2 was 5.76415e-12.
